# Repeated COVID-19 vaccine boosters elicit variant-specific memory B cells in humans

**DOI:** 10.1101/2025.10.16.682893

**Authors:** M. Alejandra Tortorici, Kaitlin R. Sprouse, Amin Addetia, Jack T. Brown, Alex Harteloo, Anna Elias-Warren, Helen Y. Chu, David Veesler

## Abstract

The first exposure to a pathogen or an antigen profoundly impacts immune responses upon subsequent encounter with related pathogens. This immune imprinting explains that infection or vaccination with currently circulating SARS-CoV-2 variants primarily recalls cross-reactive memory B cells and antibodies induced by prior Wu spike (S) glycoprotein exposure rather than priming *de novo* responses. The magnitude and persistence of immune imprinting in mRNA vaccinated populations and the prospect to overcome it are not understood. To understand the impact of immune imprinting, we investigated memory B cell and plasma antibody responses after administration of multiple doses of XBB.1.5 and JN.1/KP.2 updated COVID-19 vaccine boosters. We found that administration of the JN.1/KP.2 booster elicited broadly neutralizing antibody responses against recently circulating SARS-CoV-2 variants that were accounted for by recall of Wu S-induced immunity. We detected an increased fraction of serum antibodies and particularly memory B cells recognizing XBB.1.5 S and KP.2 S, but not Wu S, relative to individuals who received a single XBB.1.5 booster a year prior. These findings suggest that repeated exposures to antigenically divergent S trimers contribute to progressively overcoming immune imprinting and support vaccine updates and innovation to provide continued protection against COVID-19.

**In brief:** Immune imprinting due to repeated SARS-CoV-2 Wuhan-Hu-1 spike exposures is widely observed in humans. Tortorici et al. show that the humoral immune response is dominated by recall of pre-existing Wu S-induced serum antibodies and memory B cells after administration of multiple XBB.1.5 and JN.1/KP.2 COVID-19 vaccine boosters. However, the detection of an appreciable fraction of serum antibodies and particularly memory B cells binding the updated vaccine antigens, but not Wu, suggests a path towards overcoming immune imprinting

**Highlights:**

- XBB.1.5, JN.1 and KP.2 S COVID-19 vaccine boosters elicit neutralizing antibodies against current variants.
- Serum neutralizing activity against circulating variants elicited after multiple doses of updated COVID-19 vaccine boosters derive from recall of Wu S-elicited antibodies.
- Non-neutralizing antibodies specific for the updated spike antigens were detected after multiple, updated COVID-19 boosters.

## INTRODUCTION

Severe acute respiratory syndrome coronavirus 2 (SARS-CoV-2) evolves under the selective pressure of host immunity resulting from human infection and vaccination. As a result, immune evasive variants continue to emerge with spike (S) mutations eroding polyclonal plasma neutralizing antibody titers and leading to waves of (re)infections^1–8^. This requires periodic reformulation of COVID-19 vaccines to match the S sequences of circulating SARS-CoV-2 variants and maximize their effectiveness, such as the Novavax Nuvaxovid JN.1 S (protein subunit) and the Pfizer Comirnaty and Moderna Spikevax KP.2 S (mRNA) boosters. S-directed neutralizing antibodies are a primary correlate of protection against COVID-19^9–12^, with the receptor-binding domain (RBD) as the main target of serum neutralizing activity^13–16^. It is therefore essential to assess the potency and breadth of serum antibodies upon vaccine update.

Since the emergence of the first Omicron variants late 2021^17^, it became apparent that humoral immune responses to these highly divergent SARS-CoV-2 lineages are dominated by recall of pre-existing immunity originating from repeated exposure to the ancestral Wu S glycoprotein. This phenomenon is called immune imprinting or original antigenic sin and was first observed in humans infected with influenza virus^18^. The impact of immune imprinting on vaccine-elicited antibody responses in humans after release of the JN.1 and KP.2 COVID-19 vaccine boosters remains unknown, limiting our understanding of the contributions of updated antigens to humoral immunity.

To understand the persistence of immune imprinting, we studied memory B cell and serum antibody responses in humans who received multiple Wu S vaccine doses prior to administration of the updated XBB.1.5, JN.1 and KP.2 COVID-19 vaccine boosters. We found that the JN.1 and KP.2 vaccine boosters elicited broadly neutralizing serum antibodies with activity against Wu S and currently circulating variants coming from a recall of pre-existing humoral immunity induced by prior Wu S exposure. We detected serum antibody and particularly memory B cells recognizing the XBB.1.5 and KP.2 S antigens, but not Wu S, suggesting that repeated exposures to antigenically divergent S glycoproteins through vaccine updates elicit *de novo* humoral responses and progressively overcome immune imprinting.

## RESULTS

### Updated vaccine boosters elicit broadly neutralizing antibodies against circulating SARS-CoV-2 variants

To evaluate the impact of the updated COVID-19 vaccine boosters on serum neutralizing activity, we analyzed serum samples from individuals with prior infection and COVID-19 vaccination. Samples from individuals were collected at 32-60 days (median: 35) and 87-163 days (median: 97) after vaccination with two doses of XBB.1.5 S (2x XBB.1.5 S cohort, **Figure 1A**), at 31-33 days (median: 32) and 79-123 days (median: 111) after receiving a KP.2 or a JN.1 S booster from individuals who previously received two doses of XBB.1.5 S booster (2x XBB.1.5 S + 1x KP.2/JN.1 S cohort, **Figure 1B**) and at 12-33 days (median: 22) and 79-123 days (median: 95) after receiving a KP.2 or a JN.1 S booster from individuals who previously received one dose of XBB.1.5 S booster (1x XBB.1.5 S + 1x KP.2/JN.1 S cohort, **Figure 1C**) (**Table 1**). The neutralization potency and breadth of polyclonal serum antibodies was assessed with a vesicular stomatitis virus (VSV) pseudotyped with the Wuhan-Hu-1/G614 (Wu/G614), XBB.1.5, JN.1, KP.2 or KP.3 S glycoproteins (**Figures 1 and S1**). Approximately one month following KP.2/JN.1 S booster administration, serum neutralizing activity was comparable or greater for all pseudoviruses evaluated relative to subjects that only received the XBB.1.5 S booster. Geometric mean titers (GMTs) against Wu/G614, XBB.1.5, JN.1, KP.2 or KP.3 S VSV were 2,210; 1,250, 70, 70 and 60 for vaccinees in the 2x XBB.1.5 S cohort, 4,020; 1,210; 230; 480 and 320 for the 2x XBB.1.5 S + 1x KP.2/JN.1 S cohort and 2,250; 2,780; 390, 860 and 540 for the 1x XBB.1.5 + 1x KP.2/JN.1 S cohort (**Figure 1D**). Approximately three months after booster vaccination, these trends in serum neutralizing activity remained largely similar although we observed an overall waning of neutralizing activity (**Figure 1D**), as previously described^19–21^. Our data suggest that administration of the updated KP.2/JN.1 vaccine booster elicits superior serum neutralizing activity against currently circulating variants, including mismatched pseudoviruses, than XBB.1.5 boosters, supporting vaccine updates to match circulating strains.

**Figure 1.**
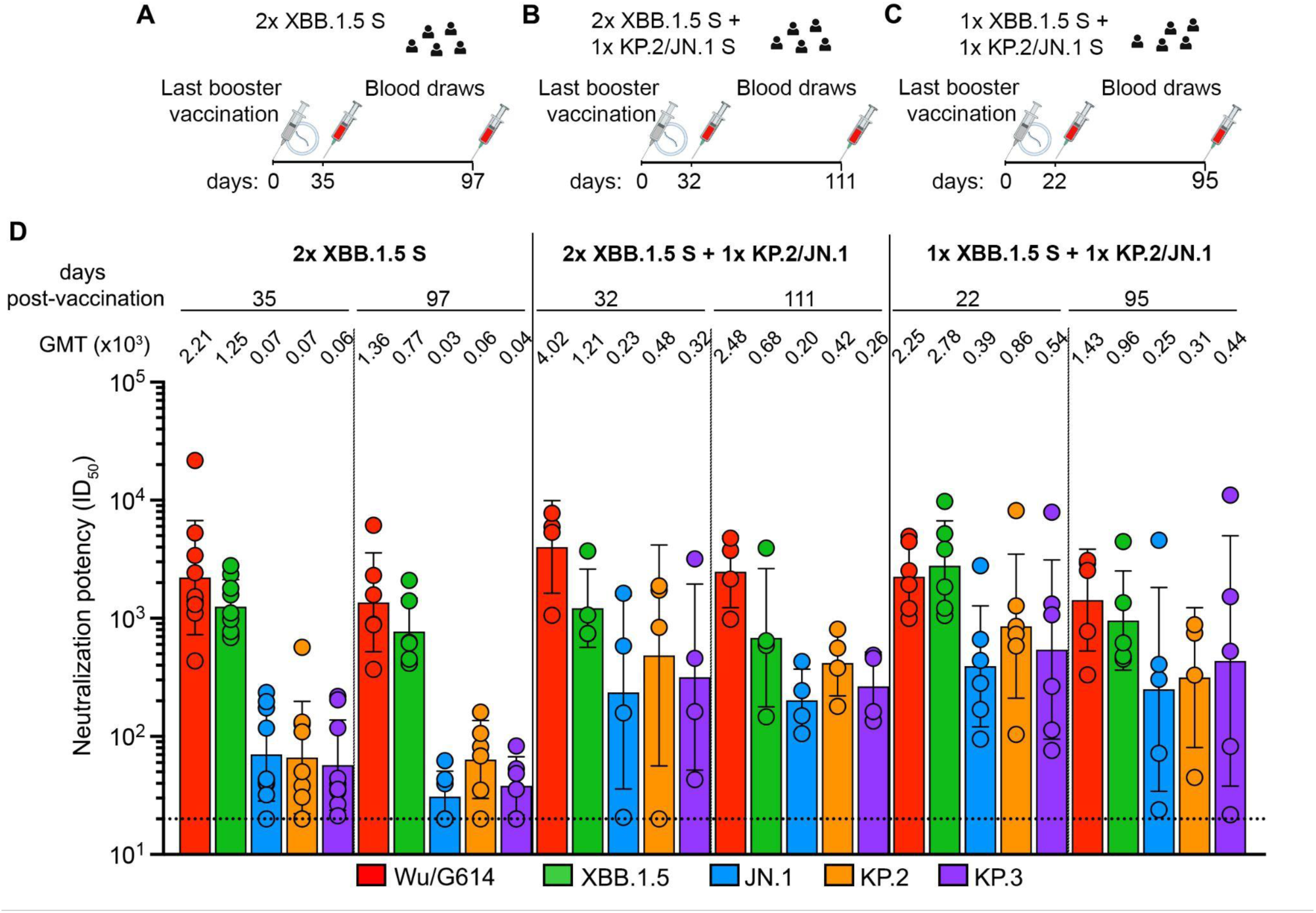
Serum neutralizing antibody responses elicited by XBB.1.5, JN.1 or KP.2 COVID-19 booster vaccination in humans. **A-C.** Timeline of vaccination and blood draws for the 2x XBB.1.5 S (A) 2x XBB.1.5 S +1x KP.2/JN.1 S (B) and 1x XBB.1.5 S + 1x KP.2/JN.1 S (C) cohorts analyzed in this manuscript. **D.** Neutralizing antibody titers evaluated using VSV pseudotyped with Wu/G614 (red), XBB.1.5 (green), JN.1 (blue), KP.2 (orange) or the KP.3 (purple) S glycoprotein using sera obtained 32-60 days (median: 35) and 87-163 days (median: 97) from 2x XBB.1.5 S cohort (A), 31-33 days (median: 32) and 79-123 days (median: 111) from 2x XBB.1.5 S + 1x KP.2/JN.1 S cohort (B) or at 12-33 days (median: 22) and 79-134 days (median: 95) from 1x XBB.1.5 S + 1x KP.2/JN.1 S cohort (C). Each data point represents the half-maximal inhibitory dilution (ID_50_) for an individual obtained from averaging two biological replicates. Geometric mean titers (GMTs) for each cohort against each pseudotyped virus are indicated above the corresponding bar graph with error bars representing the geometric SD. The horizontal dotted dashed lines indicate the limit of detection (1/20 serum dilution). See also Figure S1.

**Table 1.**
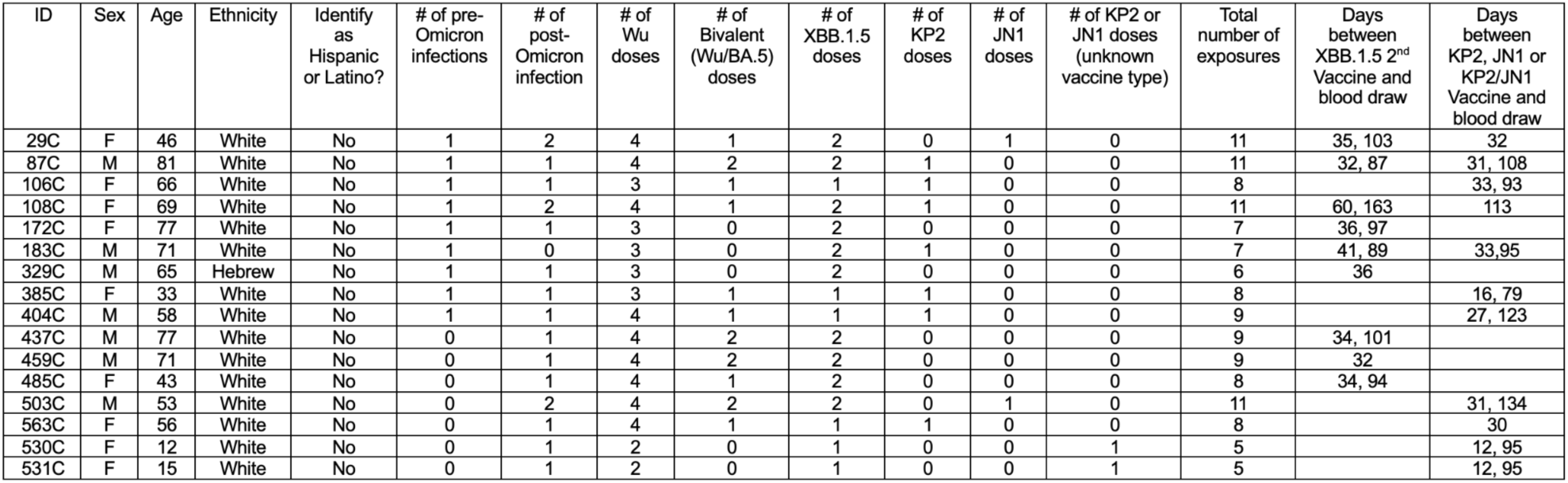
Demographics information of the cohort participants.

### Updated booster vaccination primarily recalls Wu S-directed neutralizing antibodies

The greater (or comparable) neutralization potency observed for Wu/G614 VSV relative to other pseudoviruses evaluated in all cohorts and at all time points is a serological indication of immune imprinting^22^ given that all subjects received multiple doses of updated COVID-19 boosters (lacking Wu S). We thus set out to understand the contribution of recalled Wu S-cross-reactive antibodies to the serum binding and neutralizing activity against the XBB.1.5 and KP.2 variants following vaccination with XBB.1.5 and KP.2/JN.1 booster. We depleted polyclonal serum antibodies recognizing the prefusion-stabilized Wu S and assessed remaining antibody binding titers against Wu S, XBB.1.5 S and KP.2 S ectodomain trimers using ELISAs. As expected, depletion of Wu S-directed antibodies markedly reduced Wu serum binding titers (Figures 2A **and S2**). We found that 5/9 (day 35) and 2/6 (day 97) individuals from the 2x XBB.1.5 cohort retain low but detectable levels of XBB.1.5 S-binding antibodies upon depletion (Figure 2B **and S2**). Furthermore, we observed that 4/4 (day 32) and 2/4 (day 111) individuals from the 2x XBB.1.5 S + 1x KP.2/JN.1 S cohort, and that 4/6 (day 22) and 3/5 (day 95) individuals from the 1x XBB.1.5 S + 1x KP.2/JN.1 S cohort retain detectable - albeit markedly dampened - XBB.1.5 S-binding antibodies upon depletion (Figures 2B **and S2**). Furthermore, 6/9 (day 35) and 3/6 (day 97) individuals from the 2x XBB.1.5 S cohort, 2/4 (day 32) and 3/4 (day 111) individuals from the 2x XBB.1.5 + 1x KP.2/JN.1 cohort and 4/6 (day 22) and 4/5 (day 95) retain detectable levels of KP.2-directed antibodies after depletion (Figures 2C **and S2**). Only one individual analyzed retained robust neutralization after depletion, as observed for 404C (1x XBB.1.5 S + 1x KP.2/JN.1 S cohort) inhibiting KP.2 S VSV at 22 days post vaccination (Figures 2D-F **and S1**), concurring with the high titer of KP.2-directed binding antibodies at this time point (Figures 2C **and S2**). However, no XBB.1.5 S VSV neutralization was detected post depletion for this sample despite retention of XBB.1.5-directed binding antibodies (Figure 2B). These findings show that administration of multiple updated COVID-19 boosters elicited variable magnitudes of polyclonal serum antibodies recognizing the XBB.1.5 and KP.2 S trimers without cross-reacting with Wu S. We suggest that immune imprinting is partially overcome through *de novo* elicitation of vaccine-matched antibody responses or redirection of previously existing antibody clones through affinity-maturation. However, serum neutralizing activity elicited by repeated booster administrations remains solely accounted for by a recall of Wu S cross-reactive antibodies induced by earlier exposures except for one individual (ID 404).

**Figure 2.**
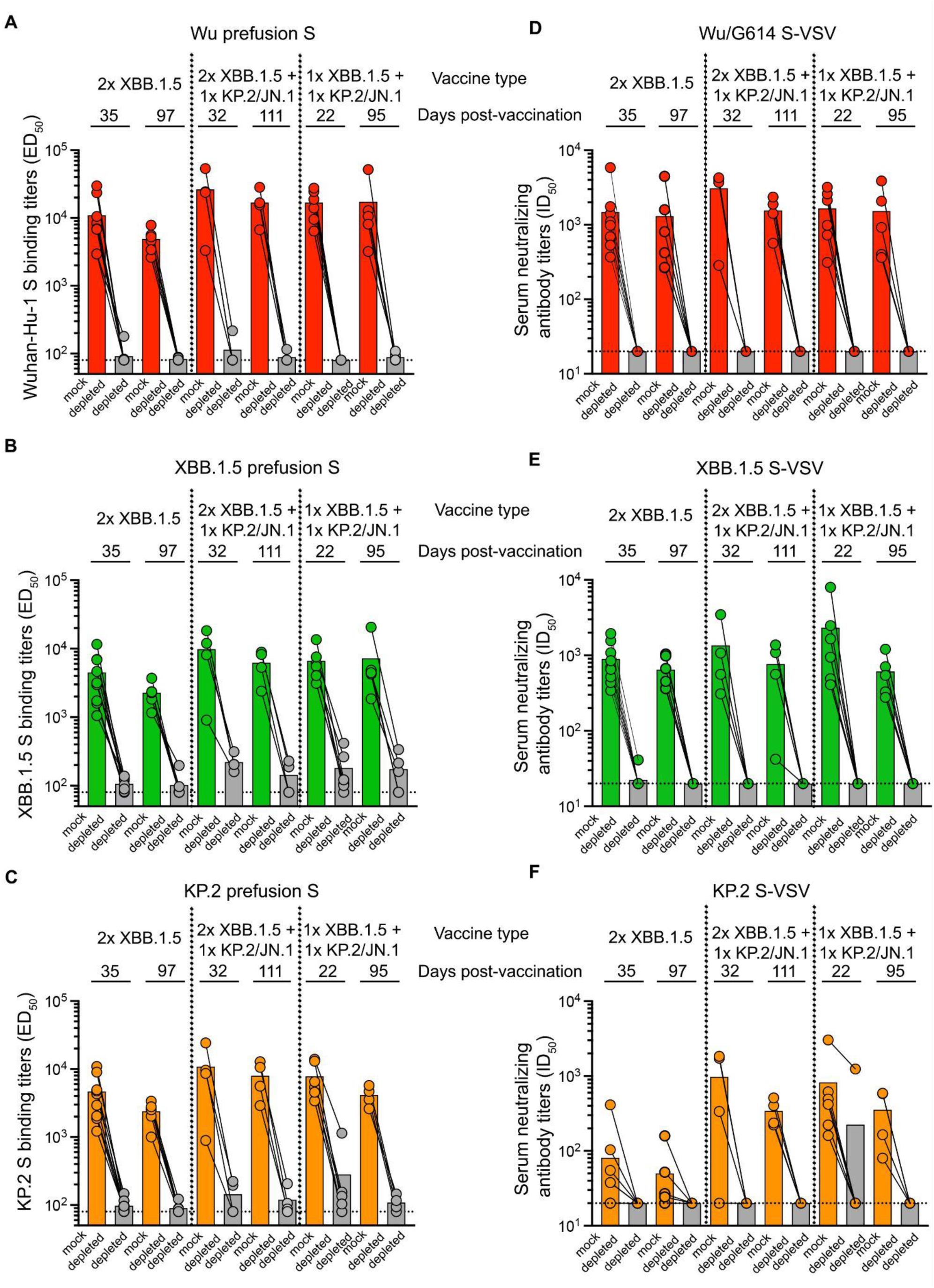
XBB.1.5 and KP.2 serum neutralizing activity is abrogated upon depletion of Wu S-reactive antibodies. (**A-C**) Antibody binding titers, expressed as half-maximal effective dilution (ED_50_), against prefusion-stabilized Wu (**A**) XBB.1.5 (**B**) and KP.2 (**C**) HexaPro S in the serum of individuals from 2x XBB.1.5 S cohort (n= 15 samples), 2x XBB.1.5 S + 1x KP.2/JN.1 S cohort (n= 8 samples) or 1x XBB.1.5 S + 1x KP.2/JN.1 cohort (n= 11 samples) boosted vaccinees determined by ELISA upon mock depletion or depletion of Wu S-reactive antibodies. (**D-F**) Neutralizing antibody titers, expressed as half-maximal inhibitory dilution (ID_50_), against Wu/G614 S VSV (**D)**, XBB.1.5 S VSV (**E**) and KP.2 S VSV (**F**) in the serum of individuals from 2x XBB.1.5 S cohort (n= 15 samples), 2x XBB.1.5 S + 1x KP.2/JN.1 S cohort (n= 8 samples) or 1x XBB.1.5 S + 1x KP.2/JN.1 cohort (n= 11 samples) boosted vaccinees upon mock-depletion or depletion of Wu S-reactive antibodies. All individuals had previously received an XBB.1.5 S mRNA booster 80-396 days (median: 251 days) before the last booster. The dotted lines indicate the limit of detection for ELISAs (1/80 serum dilution) and for neutralization assays (1/20 serum dilution). In (**A**)–(**C**), each data point represents the half-maximal effective dilution (ED_50_) for an individual obtained from the average of two biological replicates (using distinct protein batches), each comprising averaged technical duplicates. In (**D**)–(**F**), each data point represents the half-maximal inhibitory dilution (ID_50_) for an individual obtained from the average of two biological replicates (using distinct pseudovirus batches), each comprising averaged technical duplicates. Geometric mean titers (GMTs) for each cohort against each pseudovirus are shown as bar graphs. See also Figures S1 and S2.

### Repeated booster administration increases the frequency of vaccine-matched memory B cells that do not cross-react with Wu S

Prior work described large differences between the antigen-specific antibody repertoire in the memory compartment and serum antibody clonotype diversity and abundance^23–25^. To understand possible differences between these two compartments in the cohorts studied here, we analyzed memory B cell populations in the peripheral blood at the same time points as those used for analysis of serum neutralizing activity and breadth. We used flow cytometry to enumerate the frequency of memory B cells that reacted with an XBB.1.5/KP.2 S RBD pool and additionally recognized the Wu S RBD (Figure 3A). Overall, we observed that memory B cells binding to the XBB.1.5/KP.2 S RBD pool (XBB.1.5/KP.2 S RBD^+^) but not the Wu S RBD (Wu S RBD^-^), accounted for a total of 12.7 % (day 35) and 8.3 % (day 97) for the 2x XBB.1.5 cohort, 8.9 % (day 32) and 11.8 % (day 111) for the 2x XBB.1.5 + 1x KP.2/JN.1 cohort, and 31.5 % (day 22) and 29.5 % (day 95) for individuals from the 1x XBB.1.5 + 1x KP.2/JN.1 cohort (Figure 3B-D). Analysis of individual samples show that XBB.1.5/KP.2 RBD^+^ and Wu RBD^-^ memory B cells ranged in frequencies from 0 to 33.3 % (day 35) and 0 to 44.4 % (day 97) for the 2x XBB.1.5 cohort, 0-11.6 % (day 32) and 0-28.6 % (day 111) for the 2x XBB.1.5 + 1x KP.2/JN.1 cohort and from 0-51.6 % (day 22) and 15.8-36 % (day 95) for the 1x XBB.1.5 + 1x KP.2/JN.1 cohort (**Figure S3**). We note that the total number of known S exposures (vaccination and infection) is between 5-9 for the 1x XBB.1.5 + 1x KP.2/JN.1 cohort and 11 for all but one individual (for whom it is 7) for the 2x XBB.1.5 + 1x KP.2/JN.1 cohort (**Table 1**). The observation of XBB.1.5/KP.2 RBD^+^ and Wu RBD^-^ memory B cells in 7 out of 9 individuals from the 2x XBB.1.5 cohort, in 3 out of 4 from the 2x XBB.1.5 + 1x KP.2/JN.1 cohort and in 5 out of 6 individuals from the 1x XBB.1.5 + 1x KP.2/JN.1 cohort (**Figure S3**), contrasts with only 3 out of 9 subjects in a previously described cohort of subjects who received one XBB.1.5 S booster ^22^. These results suggest that immune imprinting was partially overcome (and to different extent for distinct individuals) in these cohorts through *de novo* elicitation of vaccine-matched memory B cells or redirection of previously existing clones through affinity-maturation.

**Figure 3.**
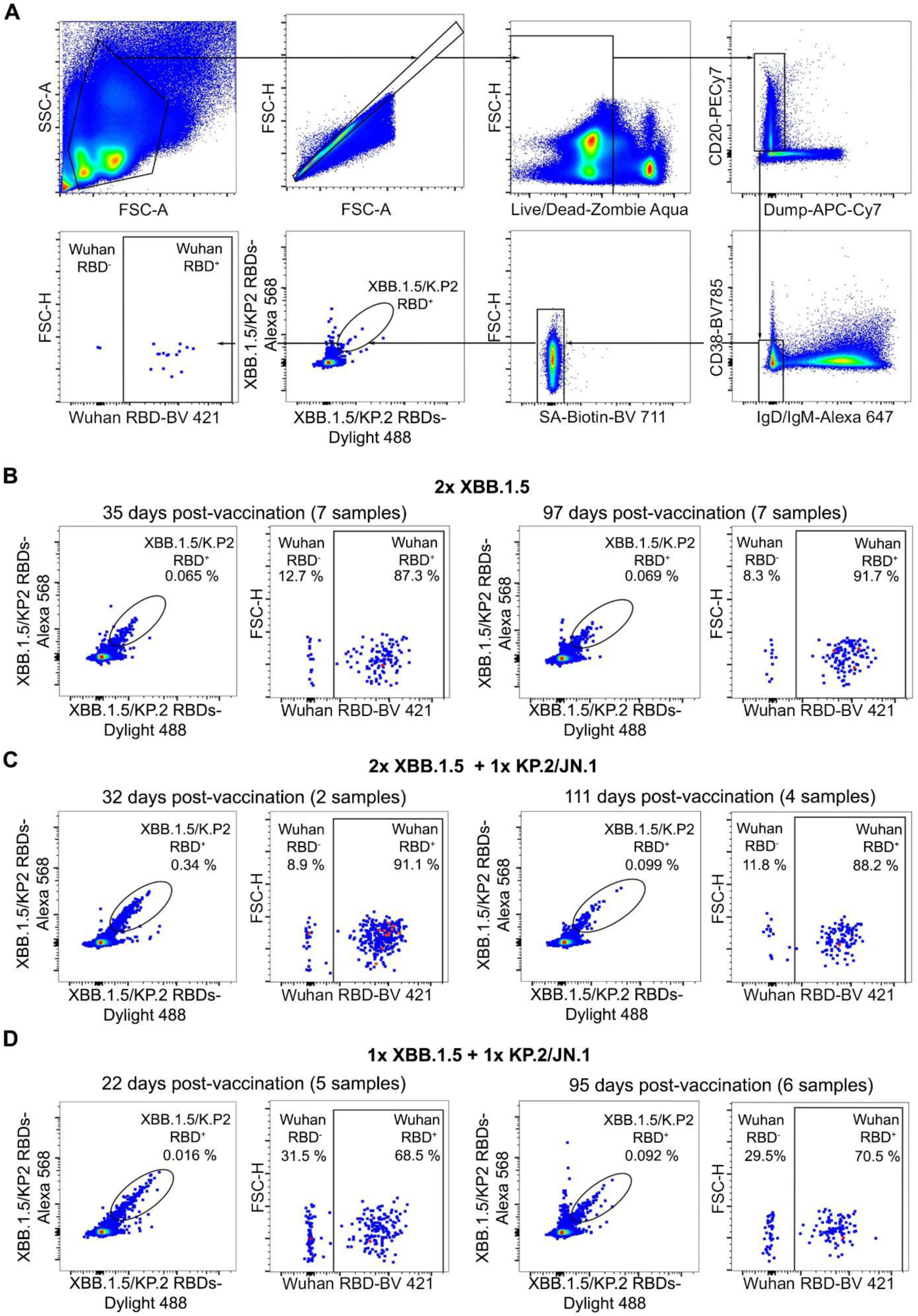
(**A**) Gating strategy to evaluate the cross-reactivity with the Wuhan-Hu-1 RBD of memory B cells binding to the XBB.1.5/KP.2 RBD pool. Dump includes markers for CD3, CD8, CD14, and CD16. The sample shown corresponds to ID: 485C, cohort 2x XBB.1.5. **(B-D)** XBB.1.5/KP.2 S RBD double-positive memory B cells (left) were analyzed for cross-reactivity with the Wuhan-Hu-1 RBD (right) using flow cytometry. Memory B cells were obtained from peripheral blood collected from individuals 32-60 (median: 35 days) or 87-163 days (median: 97 days) for the 2x XBB.1.5 cohort (**B**), 31-33 days (median: 32) and 79-123 days (median: 111) for the 2x XBB.1.5 + 1x KP.2/JN.1 cohort (**C**), and 12-33 days (median: 22 days) or 79-123 days (median: 95 days) for the 1x XBB.1.5 cohort + 1x KP.2/JN.1 cohort (**D**) after receiving the indicated vaccine booster(s). See also Figure S3.

## DISCUSSION

The ability of the human immune system to recall memory B cells elicited by prior exposure protects individuals upon subsequent encounters with the same (or closely related) pathogens. Multiple safe and effective vaccines leveraging this property were developed to provide durable protection against several human pathogens, tremendously increasing life expectancy.

Immune imprinting, also known as original antigenic sin, was first observed in studies of human immunity to influenza virus and describes how the first exposure to a virus shapes immunological outcomes of subsequent exposures to antigenically related strains^18^. Although it is often considered detrimental, imprinting during childhood resulting from exposure to seasonal influenza H1N1 or H3N2 was proposed to respectively protect against H5N1 or H7N9 zoonotic strains, most likely via hemagglutinin stem-directed antibodies^26^. This was also the case during the 2009 H1N1 “swine flu” pandemic where initial antibody responses to infection in humans were dominated by antibodies targeting the conserved hemagglutinin stem region^27,28^.

Repeated exposures through infection and vaccination during the first few years of the COVID-19 pandemic resulted in widespread Wu S imprinting. Subsequent infection with immune-evasive SARS-CoV-2 Omicron variants along with vaccine updates to match the S glycoprotein sequence of antigenically-evolved variants did not overcome immune imprinting substantially^22,25,29–38^. Instead, humoral immunity remained dominated by recall of memory B cells and plasma antibodies targeting conserved antigenic sites among SARS-CoV-2 variants, which were elicited by prior exposure.

A study of subjects vaccinated twice with inactivated SARS-CoV-2 Wu vaccines followed by repeated Omicron infections reported substantial *de novo* antibody responses elicited against these newly emerged variants^39,40^. However, these findings appear to be specific to the cohort studied, possibly due to distinct vaccine platforms and total number of exposures. Although the exact mechanisms governing Wu S imprinting remain to be elucidated, prior studies have suggested that epitope masking by pre-existing antibodies can modulate subsequent antigen-specific antibody responses^41,42^.

Here, we report that repeated administration of updated vaccine boosters to individuals with Wu S imprinting and complex exposure history elicits broadly neutralizing plasma antibody responses against matched and mismatched SARS-CoV-2 variants. Furthermore, an appreciable fraction of memory B cells recognizing the updated vaccine RBDs did not cross-react with the Wu RBD, indicating *de novo* priming of Omicron-specific naive B cells or redirection of pre-existing memory B cells leading to loss of Wu RBD binding^32,43^. We detected a higher fraction of RBD^+^ memory B cells that do not cross-react with the Wu S RBD, relative to studies carried out after prior vaccine updates^22,29,32,43^, suggesting that immune imprinting is progressively overridden. However, our depletion studies show that virtually all serum neutralizing activity is mediated by Wu S cross-reactive antibodies in all but one individual.

The rare detection of XBB.1.5 S- or KP.2 S-directed plasma antibodies that do not cross-react with Wu S contrasts with the substantially higher fraction of memory B cells binding to the XBB.1.5/KP.2 RBD pool but not the Wu RBD in most individuals. These results underscore profound differences between antibody composition and prevalence in the plasma and memory compartments, suggesting that few B cell clones give rise to long-lived bone marrow plasma cells responsible for producing plasma antibodies^23,24,44^. Collectively, these findings suggest that continued administration of updated vaccine boosters may overcome immune imprinting, concurring with the elicitation of potent *de novo* plasma neutralizing antibodies after multiple doses of updated multivalent RBD-nanoparticle vaccine doses following mRNA-1273 Wu-imprinting in non-human primates^45^. Irrespective of modulation of immune imprinting, we note that clinical evaluation of updated COVID-19 booster vaccines established their effectiveness at reducing hospitalization and severe outcomes^46–49^, motivating continued reformulation and innovation to protect the human population widely.

### Limitations of the study

Our study is based on a relatively small sample size obtained from a homogeneous population. Only a fraction of the samples could be collected longitudinally from the same individuals and the other samples were therefore obtained from different subjects at the different time points. Our analysis focused on class-switched (IgM^-^/IgD^-^) memory B cells which might have led to an underestimation of the fraction of B cells recognizing the XBB.1.5/KP.2 S RBD pool without binding to the Wu S RBD, as previously proposed ^32^.

---

**Figure S1, related to Figure 1 and 2.**
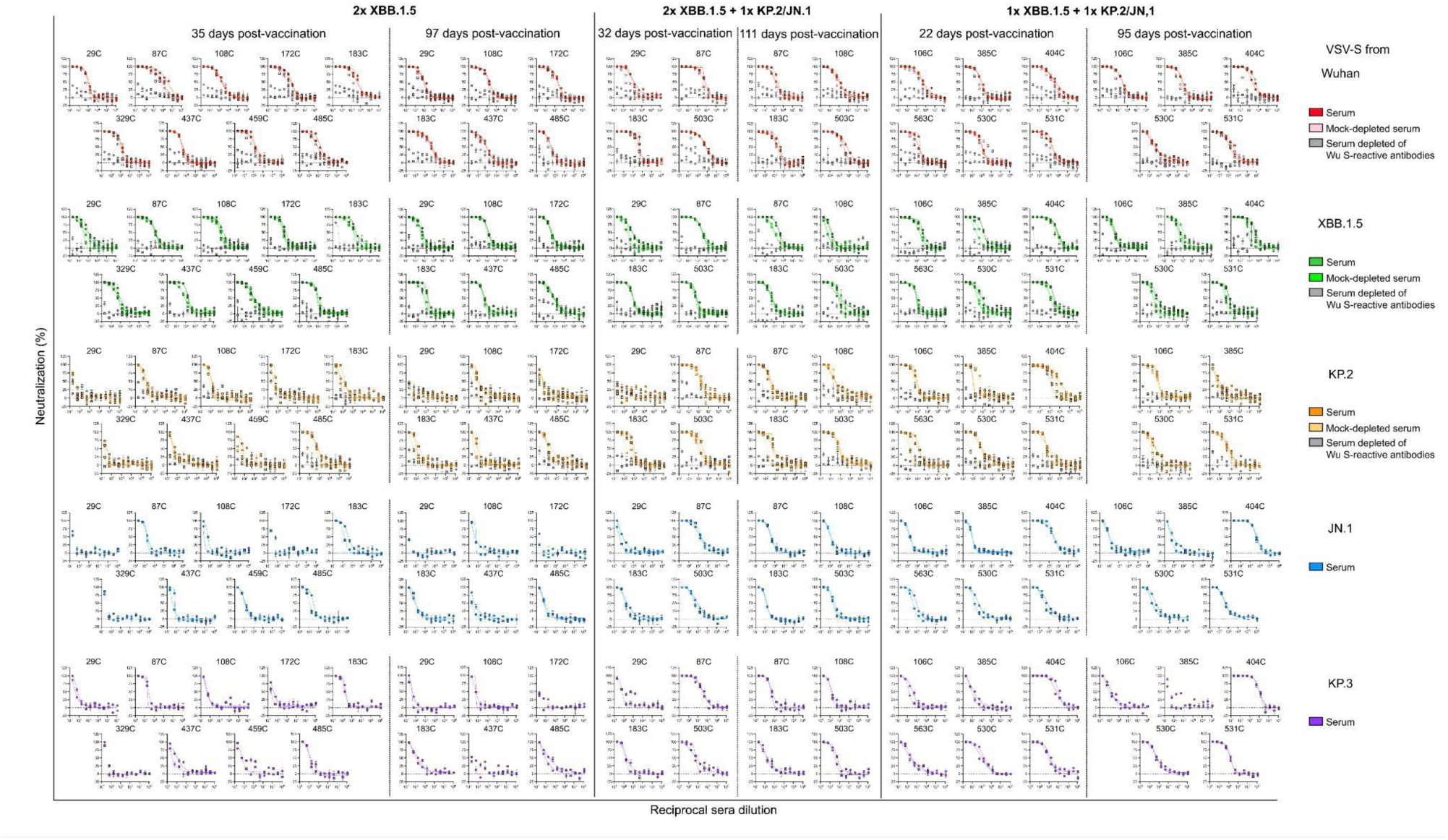
Serum neutralizing antibody titers after vaccination with XBB.1.5, JN.1 or KP.2 COVID-19 boosters. Dose-response curves against Wu/G614 S-, XBB.1.5 S, KP.2 S, JN.1 S and KP.3 S-VSV pseudoviruses mediated by human sera (Fig. 1) and by human sera that was either mock-depleted or depleted of Wu S-reactive antibodies (Fig. 2). Data is presented as the average of technical duplicates with error bars representing the SEM from two biological experiments, squares symbols/dashed line or circles symbols/solid lines. Cohort member IDs are indicated on top of each graph.

**Figure S2, related to Figure 2.**
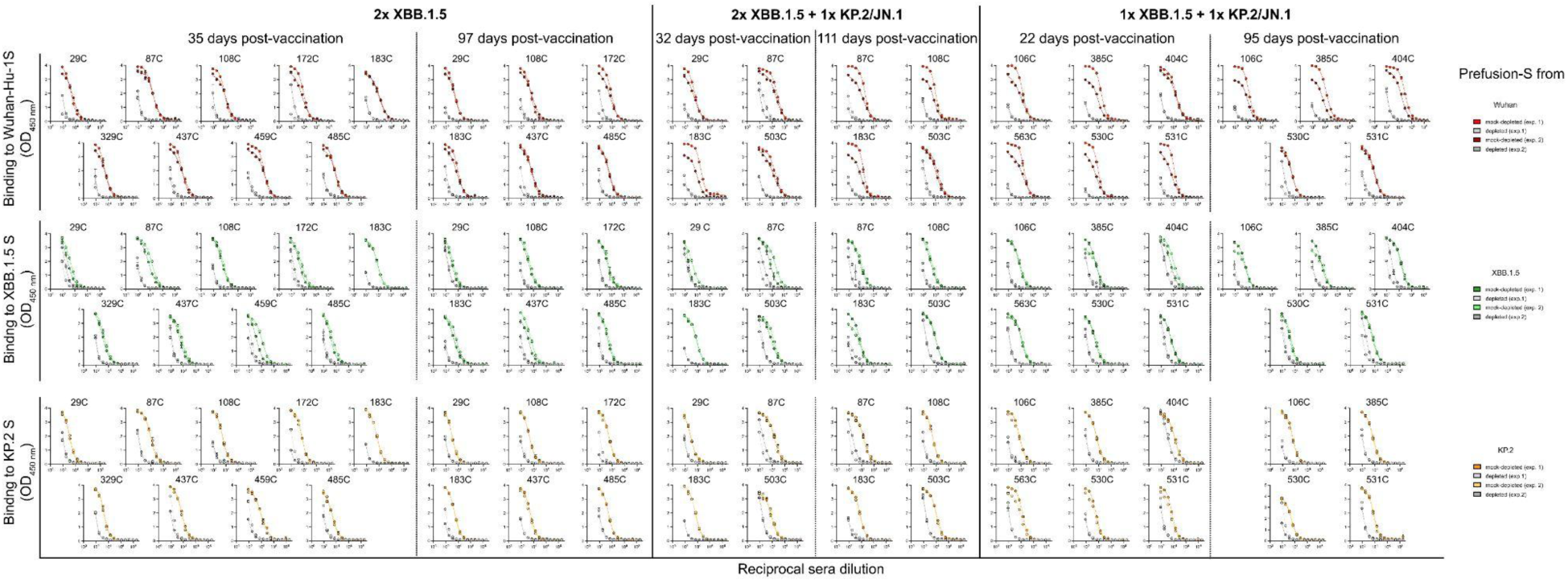
Serum antibody binding titers after vaccination with XBB.1.5, JN.1 or KP.2 COVID-19 boosters. Dose-response curves against prefusion Wu S, XBB.1.5 S and KP.2 S in human serum samples that were either mock-depleted or depleted of Wu S-reactive antibodies (grey). Data is presented as the average of technical duplicates with error bars representing the SEM from two biological experiments. Cohort member IDs are listed above each graph.

**Figure S3, related to Figure 3.**
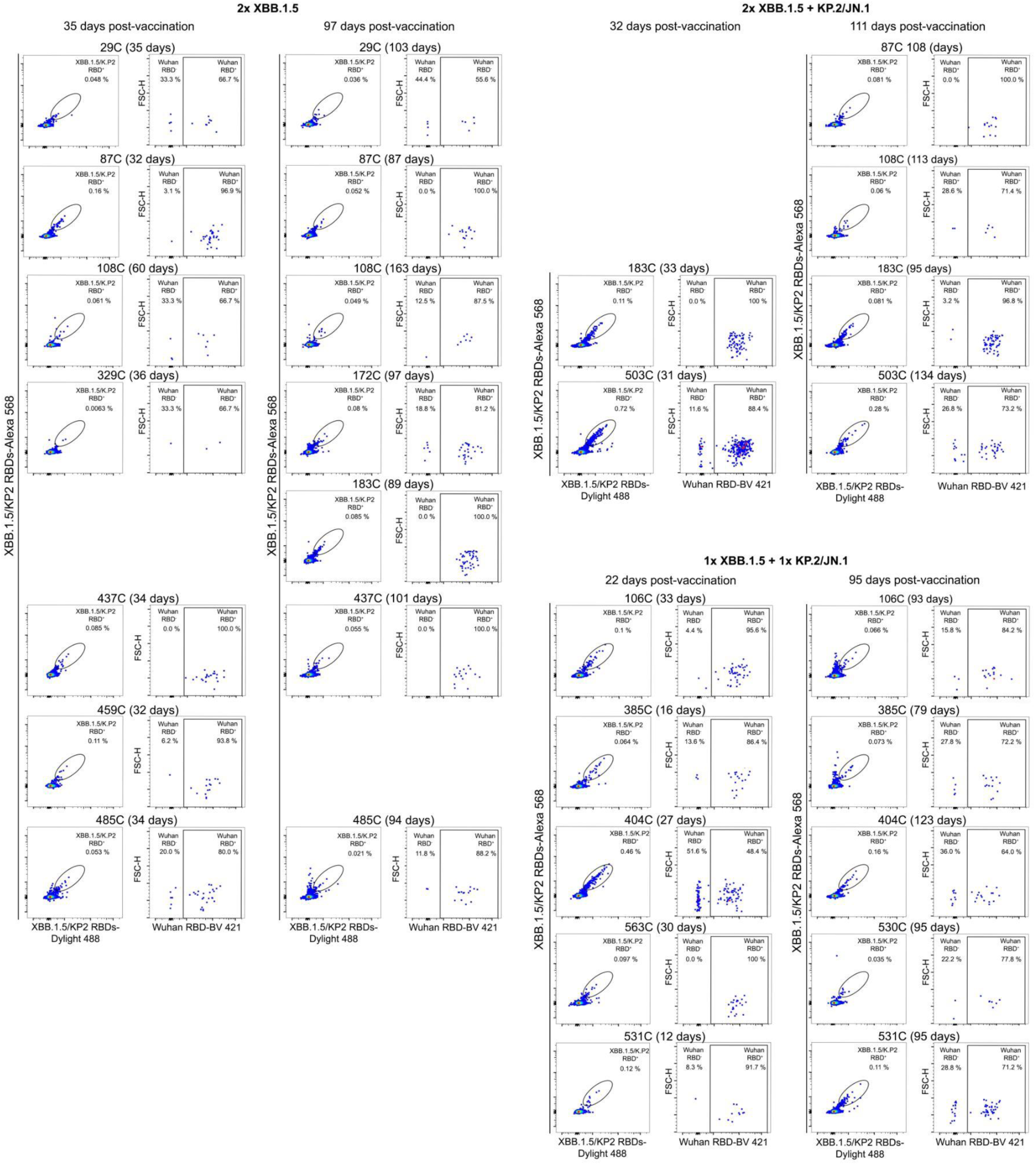
Flow cytometry analysis of memory B cells. Gating of XBB.1.5/KP.2 S RBD-reactive memory B cells and subsequent evaluation of Wu S RBD binding of these memory B cells from the peripheral blood of each individual collected at the indicated times from the 2x XBB.1.5 S (**A**), the 2x XBB.1.5 + 1x KP.2/JN.1 S (**B**) or 1x XBB.1.5 + 1x KP.2/JN.1 (**C**) cohorts using flow cytometry. The analysis was performed once.

## STAR Methods

### Key Resources table

STAR Methods Key Resources table
| REAGENT or RESOURCE | SOURCE | IDENTIFIER |
| --- | --- | --- |
| **Antibodies** |  |  |
| CD20-PECy7 | BD Biosciences | Cat# 335793; RRID:AB_399971 |
| CD3-Alexa eFluor780 | Thermo Fisher | Cat# 47-0037-41; RRID:AB_2573935 |
| CD8-Alexa eFluor780 | Thermo Fisher | Cat# 47-0086-42; RRID:AB_2573945 |
| CD14-Alexa eFluor780 | Thermo Fisher | Cat# 47-0149-42; RRID:AB_1834358 |
| CD16-Alexa eFluor780 | Thermo Fisher | Cat# 47-0168-41; RRID:AB_11219083 |
| IgM-Alexa Fluor 647 | BioLegend | Cat# 314535; RRID:AB_2566612 |
| IgD-Alexa Fluor 647 | BioLegend | Cat# 348227; RRID:AB_2563268 |
| CD38_Brilliant Violet 785 | BioLegend | Cat# 303529; RRID:AB_2561368 |
| Brilliant Violet 421 Mouse IgG | BioLegend | Cat# 400157 |
| BV11 Mouse anti-human IgG | BD OptiBuild | Cat# 740796 |
| Alexa Fluor 568 Rabbit IgG monoclonal | abcam | Cat# 1045530-17 |
| Mouse IgG1 Dylight 488 | Thermo Fisher | Cat# MA1-191-D488 |
| Goat anti-Human IgG FC Cross-Adsorb<br>Secondary Antibody, HRP | Thermo Fisher | Cat# A18823 |
| **Bacterial and virus strains** |  |  |
| *E.coli* DH10B competent cells | Invitrogen | Cat# 44-0099 |
| **Chemicals, peptides, and recombinant proteins** |  |  |
| Blocker Casein | Thermo Fisher | Cat# J600289.AP |
| Lipofectamine 2000 Transfection Reagent | Invitrogen | Cat# 11668027 |
| SureBlue Reserve TMB Microwell<br>Peroxidase Substrate, KPL | VWR | Cat# 5120-0083 |
| Flow Cytometry Staining Buffer | Thermo Fisher | Cat# 00-4222-26 |
| ExpiFectamine 293 Transfection Kit | Thermo Fisher | Cat# A14525 |
| Zombie Aqua dye | Biolegend | Cat# 423101 |
| Brilliant Stain Buffer | BD Biosciences | Cat# 563794 |
| PBS pH 7.4 | Gibco | Cat# 10010-031 |
| Penicillin-Streptomycin | Thermo Fisher | Cat# 15140163 |
| Gibco Trypsin-EDTA 0.25 % | Thermo Fisher | Cat# 25200056 |
| Fetal Bovine Serum | Thermo Fisher | Cat# S3039603HI |
| Wuhan-Hu-1 S Hexapro ectodomain | Tortorici et al. 2024 | https://www.sciencedirect.com/science/article/pii/S107476132400092X?via%3Dihub |
| XBB.1.5 S Hexapro ectodomain | Tortorici et al. 2024 | https://www.sciencedirect.com/science/article/pii/S107476132400092X?via%3Dihub |
| XBB.1.5 S RBD | Tortorici et al. 2024 | https://www.sciencedirect.com/science/article/pii/S107476132400092X?via%3Dihub |
| KP.2 S RBD | This work | N/A |
| Gibco DMEM, high glucose | Thermo Fisher | Cat# 11-965-118 |
| Opti-MEM | Thermo Fisher | Cat# 31985088 |
| Brilliant Violet 711 Streptavidin | BioLegend | Cat# 405241 |
| Brilliant Violet 421 Streptavidin | BioLegend | Cat# 405226 |
| DyLight 488 Streptavidin | BioLegend | Cat# 405218 |
| Streptavidin, Alexa Fluor 568 | Thermo Fisher | Cat# S11226 |
| **Critical commercial assays** |  |  |
| One-Glo EX Luciferase Assay System | Promega | cat# E8150 |
| BirA biotin-protein ligase standard reaction kit | Avidity | Cat# 341113 |
| **Experimental models: Cell lines** |  |  |
| HEK293T cells | ATCC | cat# CRL-11268 |
| VeroE6-TMPRSS2 (puromycin resistant) | Lempp et al. 2021 | https://www.nature.com/articles/s41586-021-03925-1 |
| Expi293F cells | Thermo Fisher | cat# A14527 |
| Il-mouse hybridoma | ATCC | CRL-2700 |
| **Recombinant DNA** |  |  |
| VSV (G*ΔG-luciferase) DNA | Kaname et al | https://journals.asm.org/doi/10.1128/jvi.02519-09 |
| pcDNA3.1(-): Wu S Hexapro ectodomain | Tortorici et al. 2024 | https://www.sciencedirect.com/science/article/pii/S107476132400092X?via%3Dihub |
| pcDNA3.1(+): XBB.1.5 S Hexapro ectodomain | Tortorici et al. 2024 | https://www.sciencedirect.com/science/article/pii/S107476132400092X?via%3Dihub |
| HDM vector: Wu/G614 S Full-length | Bowen et al. 2022 | https://www.science.org/doi/10.1126/science.abq0203 |
| HDM vector: XBB.1.5 S Full-length | Tortorici et al. 2024 | https://www.sciencedirect.com/science/article/pii/S107476132400092X?via%3Dihub |
| HDM vector: JN.1 S Full-length | Tortorici et al. 2024 | https://www.sciencedirect.com/science/article/pii/S107476132400092X?via%3Dihub |
| pcDNA3.1(+): KP.2 S Full-length | GeneScript | This study |
| HDM vector: KP.3 S Full-length | GeneScript | This study |
| pcDNA3.1 (-): Wu S RBD | Walls et al. 2020 | https://www.sciencedirect.com/science/article/pii/S0092867420302622?via%3Dihub |
| pcDNA3.1(+): XBB.1.5 S RBD | Addetia et al. 2023 | https://www.nature.com/articles/s41586-023-06487-6 |
| pCDNA3.1(+): KP.2 S RBD | GeneScript | This study. |
| **Software and algorithms** |  |  |
| FlowJo 10.8.1 | BD Biosciences | https://www.flowjo.com/ |
| BioRender | N/A | https://www.biorender.com/ |
| Prism 10 | GraphPad | https://www.graphpad.com/ |
| **Other** |  |  |
| Ultracomp eBeads plus | Thermo Fisher | Cat# 01-3333-42 |
| ArC reactive beads | Thermo Fisher | Cat# A10346 A |
| ArC negative beads | Thermo Fisher | Cat# A10346 B |
| poly-D-lysine-coated 10 cm dishes | Thermo Fisher | Cat# 08-757-466 |
| Immuno Nonsterile 384-well plates | Thermo Fisher | Cat# 464718 |
| Costar Assay Plate, 96 well white with clear flat bottom | Corning | Cat# 3903 |
| EndoFree Plasmid Mega kit | Qiagen | Cat# 12381 |
| His-Tag Dynabeads | Thermo Fisher | Cat# 10104D |
| DynaMag-2 Magnet | Thermo Fisher | Cat# 12-321-D |
| 100-kDa cut-off concentrator | Amicon | Cat# UFC910024 |
| 30-kDa cut-off concentrator | Amicon | Cat# UFC903096 |
| 10-kDa cut-off concentrator | Amicon | Cat# UFC901096 |
| Sepharose excel histidine-tagged protein purification resin | Cytiva | Cat# 17371203 |
| Superdex 200 Increase 10/300 GL | Cytiva | Cat# 28990944 |
| Polystyrène round-bottom tube with cell-strainer cap | Falcon, a corning brand | Cat# 352235 |

## CONTACT FOR REAGENT AND RESOURCE SHARING

### Lead Contact

Further information and requests for resources and reagents should be directed to and will be fulfilled by the lead contact David Veesler.

### Materials availability

Requests for reagents and resources generated in this study will be fulfilled by the lead contact.

Data and code availability

- All the data are presented in this manuscript.
- This manuscript does not report original code.
- Any additional information required to reanalyze the data reported here, is available from the lead contact upon request.

## ACKNOWLEDGMENTS

This study was supported by the National Institute of Allergy and Infectious Diseases (DP1AI158186, P01AI167966, and 75N93022C00036 to D.V.); a Pew Biomedical Scholars Award (to D.V.); an Investigators in the Pathogenesis of Infectious Disease Awards from the Burroughs Wellcome Fund (to D.V.); Fast Grants (to D.V.); the University of Washington Arnold and Mabel Beckman Cryo-EM Center; and the National Institute of Health grant S10OD032290 (to D.V.). D.V. is an investigator of the Howard Hughes Medical Institute and the Hans Neurath Endowed Chair in Biochemistry at the University of Washington.

## AUTHOR CONTRIBUTIONS

M.A.T., A.A., and D.V. designed the study. M.A.T. and K.S. produced pseudoviruses. M.A.T. performed serum depletion experiments. M.A.T and A.A. carried out flow cytometry experiments. K.S. carried out neutralizations and ELISAs assays. K.S., J.B. and C.S. recombinantly produced proteins. A.H. and A.E.W. recruited the cohort’s members and prepared sera and PBMCs under the supervision of H.Y.C. M.A.T. and D.V. analyzed the data and wrote the manuscript, with input of all the authors.

## EXPERIMENTAL MODEL AND STUDY PARTICIPANT DETAILS

### Cell lines

VeroE6-TMPRSS2 (puromycin resistant) is a female cell line generated using a lentivirus system. Expi293F is a female cell line (obtained from Thermo Fisher). HEK293T is a female cell line (obtained from ATCC) and Il-mouse hybridoma is a female cell line (obtained from ATCC). Cells were cultivated at 37°C in an atmosphere of 5 % CO_2_ for adherent cells and 8 % CO_2_ with 130 rpm of agitation for suspension cells. None of the cell lines used were routinely tested for mycoplasma contamination.

### Sample donors and collection

Sera and PBMCs were collected after informed consent from participants in the prospective longitudinal Hospitalized or Ambulatory Adults with Respiratory Viral Infections (HAARVI) study from Washington State, USA, which was approved by University of Washington Institutional Review Board (protocol #STUDY00000959). Demographics information is provided in Table 1.

## METHOD DETAILS

### Plasmids

All genes used in this study were synthesized by GenScript, inserted in frame with a Kozak sequence to direct translation and codon optimized for expression in mammalian cells. Plasmid encoding for Wu S Hexapro ectodomain (residues 1-1208) and XBB.1.5 S Hexapro (residues 1-1203) were previously described in Tortorici et al., 2024. Plasmid encoding KP.2 S Hexapro ectodomain (residues 1-1204) was cloned into pcDNA 3.1 (+) and, like constructs Wu S Hexapro and XBB.1.5 S Hexapro, harbors the HexaPro mutations (Hsieh et al., 2020), a wildtype signal peptide, a furin cleavage site mutated _685_RSV_687_ to _685_SSV_687_, an avi-tag and an octa-his tag for affinity purification.

Constructs for membrane-anchored S glycoproteins for SARS-CoV-2 Wu/G614 and XBB.1.5 were previously described^22^. Membrane-anchored S glycoprotein constructs for JN.1 (1-1248), KP.2 (1-1248) and KP.3 (1-1248) were all cloned into an HDM vector with the authentic signal peptide and harboring a C-terminal deletion of 21 residues^51^. Mutations included in the variants XBB.1.5, JN.1, KP.2 and KP.3 are described in Table 2.

**Table 2.** Mutations in the SARS-CoV-2 variants XBB.1.5, JN.1, KP.2 and KP.3 S full-length relative to Wuhan-Hu-1/G614 S full-length. Ectodomains of the same variant genes contain in addition the hexapro mutations ^50^. Residue numbering is based on Wuhan-Hu-1/G614 S.

| Variant | Mutations |
| --- | --- |
| XBB.1.5 | T19I, L24del, P25del, P26del, A27S, V83A, G142D, Y144del, H146Q, Q183E, V213E, G252V, G339H, R346T, L368I, S371F, S373P, S375F, T376A, D405N, R408S, K417N, N440K, V445P, G446S, N460K, S477N, T478K, E484A, F486P, F490S, Q498R, N501Y, Y505H, H655Y, N679K, P681H, N764K, D796Y, Q954H, N969K |
| JN.1 | 17M, L18P, T19L, F20ins, T20N, R21L, T22I, Q23T, L24T, P25T, P26Q, A27S, S50L, H69del, V70del, V127F, G142D, Y144del, F157S, R158G, N211del, L212I, V213G, L216F, H245N, A264D, I332V, G339H, K356T, S371F, S373P, S375F, T376A, R403K, D405N, R408S, K417N, N440K, V445H, G446S, N450D, L452W, L455S, N460K, S477N, T478K, N481K, V483del, E484K, F486P, Q498R, N501Y, Y505H, E554K, A570V, P621S, H655Y, I670V, N679K, P681R, N764K, D796Y, S939F, Q954H, N969K, P1143L |
| KP.2 | JN.1 mutations plus F456L, R346T, V1104L |
| KP.3 | JN.1 mutations plus F456L, Q493E, V1104L |

### Recombinant protein production

To express Wu, KP.2 and XBB.1.5 stabilized S ectodomains with the hexapro mutations, Expi293 cells were transfected with the corresponding plasmids using Expifectamine following the manufacturer’s instructions. Five days post-transfection, supernatants were clarified by centrifugation at 4,121 x g for 30 minutes, supplemented with 25 mM sodium phosphate pH 8.0, 300 mM NaCl. Supernatant was vacuum filtered (0.22 µm) and mixed with Ni Sepharose Excel resin (Cytiva), previously equilibrated in 25 mM sodium phosphate pH 8.0, 300 mM NaCl. After 2 h incubation at room temperature in a roller shaker, supernatant was added to a gravity flow column and washed with 25 mM sodium phosphate pH 8.0, 300 mM NaCl and 50 mM imidazole. Proteins were eluted with 25 mM sodium phosphate pH 8.0, 300 mM NaCl and 500 mM imidazole and buffer exchanged in 25 mM sodium phosphate pH 8.0, 150 mM NaCl using a centrifugal device (Amicon) with a MWCO of 100 kDa and stored at 4°C or immediately used.

### Sera antibody depletion

Invitrogen His-Tag Dynabeads (Thermo Fisher) were used for depletion of serum samples from antibodies recognizing the Wu S trimer, as previously described^22^ with some modifications. Vortexed beads were incubated at room temperature on a DynaMag-2 Magnet (Thermo Fisher) for 2 min to allow beads to separate for the liquid phase. Supernatant was discarded and beads were washed one time with TBS-T (20 mM Tris-HCl pH7.5, 150 mM NaCl, 0.1 % (w/v) Tween 20) and divided in two tubes. After a 2 min incubation on the magnet, TBS-T supernatants were discarded and one set of beads was incubated with 4 mg of purified his-tagged Wu S ectodomain trimer in purification buffer (Wu S depletion) and the other set was incubated with TBS-T alone (mock depletion) with gentle rotation for 1 h at room temperature. Supernatants were discarded using the magnet and beads were washed three times with TBS-T. Subsequently, 20 ml of each of the sera samples were incubated with 80 ml of S-loaded beads or mock-loaded beads for 1 h at 37°C during which time they were mixed every 15 min. Sera samples were separated from the beads using the magnet and subjected to a second round of depletion using the same protocol as described for the first round. Incubated supernatants were recovered using a magnet to separate them from the beads and used for neutralization and binding assays.

### VSV pseudotyped virus production and neutralization

Vesicular stomatitis virus (VSV) was pseudotyped with SARS-CoV-2 Wu S harboring the G614, XBB.1.5, KP.2, KP.3 or JN.1 mutations following a previously described protocol (Tortorici et al., 2024). Briefly, HEK293T cells seeded in poly-D-lysine-coated 10-cm dishes in DMEM (Gibco, Thermo Fisher) supplemented with 10 % FBS and 1 % PenStrep were transfected with 24 µg of the corresponding plasmid, 60 µl Lipofectamine 2000 (Life Technologies) in 3 ml of Opti-MEM (Thermo Fisher), following the manufacturer’s instructions. Transfection mixture was added to 10 cm plates containing washed cells and 7mL of DMEM and 10% FBS. The next day, cells were washed three times with DMEM and were transduced with VSV `Δ`G-luc.79 After 2 h, the virus inoculum was removed and cells were washed five times with DMEM prior to the addition of DMEM supplemented with anti-VSV-G antibody [Il-mouse hybridoma supernatant diluted 1 to 25 (v/v), from CRL-2700, ATCC] to minimize parental background. After 18-24 h, supernatants containing pseudotyped VSV were harvested, centrifuged at 2,000 x g for 10 min to remove cellular debris, filtered with a 0.45 mm membrane, concentrated 10 times using a 30 kDa cut-off membrane (Amicon), aliquoted, and frozen at −80°C.

For VSV pseudotyped neutralizations, VeroE6-TMPRSS2 cells were seeded into coated clear bottom white walled 96-well plates at 20,000 cells/well and cultured overnight at 37°C. Twelve 3-fold serial dilutions of each sera sample were prepared in DMEM. Pseudotyped VSV viruses, diluted 1 to 10 in DMEM containing anti-VSV-G antibody [Il-mouse hybridoma supernatant diluted 1 to 25 (v/v), from CRL-2700, ATCC], were added 1:1 (v/v) to each sera sample dilution and mixtures of 40 µl volume were incubated for 45-60 min at room temperature. VeroE6-TMPRSS2 cells were washed three times with DMEM and the mixtures pseudovirus/ sera dilutions were added to the cells. Two hours later, 40 µl of DMEM were added to the cells. After 17-20 h, 40 µl of One-Glo-EX substrate (Promega) were added to each well and incubated on a plate shaker in the dark at 37°C. After 5-15 min incubation, plates were read on a Biotek Neo2 plate reader. Measurements were done in duplicate with at least two biological replicates. Relative luciferase units were plotted and normalized in Prism 10 (GraphPad): cells without pseudotyped virus added were defined as 0 % infection or 100 % neutralization, and cells with virus only (no sera) were defined as 100 % infection or 0 % neutralization.

### Enzyme-linked immunosorbent assays (ELISA)

Analysis of sera binding antibodies for samples mock-depleted or depleted of antibodies binding to the Wu S ectodomain trimer was performed using ELISAs. Briefly, clear flat bottom Immuno Nonsterile 384-well plates (Thermo Fisher) were coated overnight at room temperature with 30 µl of Wu S, XBB.1.5 S or KP.2 S prepared at 3 ng/ml per well in PBS (137 mM of NaCl, 2.7 mM of KCl, 10 mM of Na_2_HPO4, and 1.8 mM of KH_2_PO_4_, pH 7.2). The next day, plates were blocked with Blocker Casein (Thermo Fisher) at 37 C and subsequently incubated with serial dilutions of sera samples for 1 h at 37°C. After four washing steps with TBS-T, goat anti-human IgG-Fc secondary antibody conjugated to HRP (Thermo Fisher), diluted 1/500 was added and incubated for 1 h at 37°C. Plates were washed four times with TBS-T and KPL SureBlue Reserve TMB Microwell Peroxidase Substrate (VWR) was added. After 2 min incubation, 1N HCl was added and absorbance at 405 nm was measured using a Biotek Neo2 plate reader. Data were plotted using Prism 10 (GraphPad).

### Flow cytometry analysis of SARS-CoV-2 RBD-reactive memory B cells

To define specific B cell populations reactive with the XBB.1.5 or KP.2 and Wu RBDs, RBD–streptavidin tetramers conjugated to fluorophores were generated by incubating the corresponding biotinylated RBDs with streptavidin at a 4:1 molar ratio for 30 min at 4°C. Excess of free biotin was added to the reaction to bind any unconjugated sites in the streptavidin tetramers. The RBD-streptavidin tetramers were washed once with PBS (137 mM NaCl, 2.7 mM KCl, 10 mM Na_2_HPO4, 1.8 mM KH_2_PO4, pH 7.4) and concentrated with a 100-kDa cut-off centrifugal concentrator (Amicon). An additional streptavidin tetramer conjugated to biotin only was generated and included in the staining as decoy.

Approximately 5 to 10 million PMBCs were collected 7-13 days and 30-63 days post-vaccination for individuals who received an XBB.1.5 S mRNA vaccine booster. Cells were collected by centrifugation at 1000 g for 5 min at 4°C, washed twice with PBS and stained with Zombie Aqua dye (Biolegend) diluted 1:100 in PBS at room temperature. After 30 min incubation, cells were washed twice with PBS and stained with antibodies for CD20-PECy7 (BD), CD3-Alexa eFluor780 (Thermo Fisher), CD8-Alexa eFluor780 (Thermo Fisher), CD14-Alexa eFluor780 (Thermo Fisher), CD16-Alexa eFluor780 (Thermo Fisher), IgM-Alexa Fluor 647 (BioLegend), IgD-Alexa Fluor 647 (BioLegend), and CD38-Brilliant Violet 785 (BioLegend), all diluted 1:200 in Brilliant Stain Buffer BD), along with the RBD-streptavidin tetramers for 30 min at 4°C. Cells were washed three times, resuspended in PBS, and passed through a 5 ml Polystyrene round-bottom tube with cell-strainer cap (Falcon) before being examined on a BD FACSymphony A3 for acquisition and FlowJo 10.8.1 for analysis. Gates for identifying the XBB.1.5 or KP.2 RBD double-positive population as well as the subsequent Wu RBD-positive and Wu RBD-negative populations were drawn based on staining of fluorescent minus one as controls.

